# Integrative Multi-Omics Analysis of Gut Microbiota Dysbiosis and Host–Microbiome Interaction Mechanisms in Hypertension

**DOI:** 10.64898/2026.06.18.733290

**Authors:** Wenkai Lai, Shaoping Huang, Yuchen Zhang, Shirong Lai, Shanwen Sun, Furong Tang, Haidan Yan, Fenglong Yang

## Abstract

**Objective:** To characterize gut microbiota dysbiosis in hypertension and investigate its multilevel interactions with the host immune system.

**Methods:** Integrated multi-cohort microbiome data were used to evaluate microbial diversity, differential abundance, and co-occurrence network features between individuals with hypertension and healthy controls. The scBPS framework was applied to analyze microbiome–cell associations, enabling the resolution of relationships between key microbial taxa and functional states of immune cells at single-cell resolution.

**Results:** Several potentially protective genera reduced in hypertension and occupied central topological positions in the co-occurrence networks. Single-cell analyses further demonstrated that multiple key genera were closely associated with the functional states of monocytes and T cells (p<0.05). Specifically, *Bacteroides* and *Bifidobacterium* were associated with the proliferation and repair of classical monocytes; *Butyricimonas* showed a negative association with antigen processing and presentation pathways in monocytes; and *Oscillospira* promoted the transition of dnT cells toward an immunoregulatory state, suggesting its potential role in immune homeostasis.

**Conclusions:** Integrated multi-omics analyses reveal that hypertension-associated gut microbes may contribute to disease development through immune regulation, providing insights into microbiome–immune interaction mechanisms and potential targets for precision interventions.

## Introduction

Hypertension remains one of the major risk factors for cardiovascular diseases (CVD) such as stroke and heart failure. In addition, it is an important associated factor for common comorbidities such as chronic kidney disease, obesity, and type 2 diabetes [1, 2]. In 2010, approximately 31% of the global population had hypertension, making it a global public health issue[3]. Despite extensive research and interventions, blood pressure control still faces many challenges[4].

The gastrointestinal tract harbors the largest population of immune cells in the body and represents a critical interface between the environment and the host[5]. Lifestyle can shape and be influenced by the microbiome[6, 7, 8], thereby altering the risk of developing hypertension. Emerging evidence indicates that the gut microbiota plays a pivotal role in the development of cardiovascular diseases[9]. For instance, patients with heart failure with preserved ejection fraction (HFpEF) exhibit pronounced gut microbial dysbiosis, with microbial profiles that differ substantially from those of healthy individuals[10]. The homeostasis of the gut microbiota is considered fundamental to human health; however, disruption of this balance leads to gut dysbiosis[11]. Certain gut bacteria can influence vascular health and blood pressure regulation through the metabolism of bioactive compounds such as short-chain fatty acids (SCFAs), whereas microbial dysbiosis may increase the risk of cardiovascular diseases[12].

The gut microbiota is not only involved in digestion and metabolic processes but also plays essential roles in immune regulation, neuro-transmission, and endocrine signaling. Through dynamic interactions with the host, the gut microbiome can profoundly influence host metabolic status, immune responses, and the ability to adapt to environmental stimuli, thereby playing a critical role in maintaining host health[13]. However, impairment of gut immune regulation may lead to local inflammation, subsequently compromising intestinal barrier integrity and facilitating the translocation of immunogenic substances such as lipopolysaccharide (LPS) from the gastrointestinal tract into the peripheral circulation[14]. These highly stimulatory molecules can trigger pro-inflammatory responses beyond the gut. Accumulating evidence indicates that chronic pro-inflammatory processes are closely associated with elevated blood pressure and represent a hallmark of cardiovascular diseases[15]. In this context, immune cells—including macrophages, dendritic cells, natural killer (NK) cells, B lymphocytes, and T lymphocytes—play pivotal roles[16]. Following infiltration into target organs, immune cells secrete cytokines (such as interleukins, interferons, and tumor necrosis factors[17]) that modulate inflammation, oxidative stress, and renal sodium and water retention, ultimately exacerbating functional impairment, remodeling, and fibrosis of the vasculature, kidneys, and heart[18]. Interactions between immune cells and the vascular wall, including the release of cytokines such as TNF-α[19], interleukin-17A[20], and interferon-γ[21], contribute to increased systemic vascular resistance and the generation of reactive oxygen species (ROS)[19].

A more specific example is that *Akkermansia muciniphila* can induce the production of IgG1 antibodies in mice and elicit antigen-specific T-cell responses[22]. Microbial antigens derived from *A. muciniphila* can also induce immune tolerance and promote the differentiation of naive *CD*4^+^*CD*44^−^*Foxp*3^−^ T (Tn) cells into the Treg lineage[23], thereby regulating host inflammatory responses. Collectively, the human gut microbiota plays a key role in maintaining tissue homeostasis and is associated with a wide range of diseases affecting multiple organs[24, 25, 26]. Evidence from animal models[27] and human cohorts[28] indicates that certain gut microbes exhibit significant heritability, motivating efforts to identify host genomic risk loci associated with the gut microbiota through genome-wide association studies (GWAS)[29].

However, the heterogeneity of cellular composition within organs limits our understanding of host–microbiome interactions at the cellular level. In particular, most hypertension-related studies to date have primarily focused on analyses of the microbiome and metabolome, and there is still a lack of research directly linking gut microbiota to interactions with specific host cell types. Recently, scBPS (single-cell Bacterial Polygenic Score)[30], a software framework specifically designed to integrate microbial genome-wide association studies with single-cell transcriptomics, has been developed. This approach enables the investigation of interaction-associated relationships between gut microbiota and host cell populations at single-cell resolution.

Accordingly, this study aims to integrate multi-omics data to perform differential genera analysis across multiple independent gut microbiome cohorts, thereby systematically identifying microbial genera strongly associated with hypertension. Building upon these findings, we applied the scBPS framework to investigate the interactions between gut microbiota and distinct subpopulations of host peripheral blood mononuclear cells (PBMCs) at single-cell resolution. Through this integrative approach, we sought to elucidate the regulatory interplay between hypertension-associated gut microbes and host immune cell subsets and their collective impact on blood pressure homeostasis. In addition, causal machine learning approaches were employed to perform causal inference on the observed statistical associations, thereby strengthening the interpretability of the results from a causal perspective. Our analyses revealed that *Bacteroides* and *Bifidobacterium* are closely associated with immune reprogramming of classical monocytes, *Butyricimonas* plays a critical role in antigen presentation by classical monocytes, and *Oscillospira* is strongly linked to the immunosuppressive functions and inflammatory responses of double-negative T (dnT) cells. By integrating multi-omics data, this study provides novel insights into the complex interactions among gut microbiota and host immune cells, particularly highlighting their dysregulated interplay in hypertension. Collectively, these findings not only advance our understanding of microbiota–metabolite imbalances and their immune-mediated effects in hypertension but also offer a theoretical foundation for the development of microbiota-targeted intervention strategies.

## Materials and methods

### Data Sources and Preprocessing

As summarized in Table 1, this study integrated multiple gut microbiome cohorts from diverse sources. Specifically, a total of 5349 samples were obtained from the Chinese Prospective Hypertension 16S Cohort (CPH16S), which comprises 16 prospective Chinese hypertension cohorts reported by Zhou HW et al.[31]. In addition, 441 samples were collected from the Colombian Hypertension 16S Dataset (CH16S) reported by Jacobo de la Cuesta-Zuluaga et al.[32]. Furthermore, 218 samples from a Chinese hypertension metagenomic dataset (Chinese Hypertension Metagenomic Dataset, CHMG) were retrieved from the curatedMetagenomicData database. Collectively, these cohorts cover a broad range of ages and body mass index (BMI). For quality control, samples from individuals who had received antihypertensive treatment or had a history of antibiotic use were excluded from the analysis.

**Table 1.** Summary of Gut Microbiome and Single-Cell Transcriptomic Datasets.

|  | Region (Proportion) | Data Type | Sample Size<br>(Hypertension /<br>Healthy Controls) | Data Source |
| --- | --- | --- | --- | --- |
| CPH16S | China (100%) | 16S rRNA sequencing | 5,349 (1,645 / 3,704) | [31] |
| CH16S | Colombia (100%) | 16S rRNA sequencing | 441 (110 / 331) | [32] |
| CHMG | Austria (6.5%), China (86.7%), Italy (6.9%) | Metagenomics | 218 (169 / 49) | curatedMetagenomicData database |
| Single-cell transcriptomic data | China (100%) | Single-cell RNA sequencing | 25 (12 / 13) | NCBI |

To further investigate the associations between gut microbiota and host immune cells, we additionally collected single-cell RNA sequencing data of peripheral blood mononuclear cells (PBMCs) from 25 individuals in the PRJNA1273480 project[33]. Moreover, genome-wide association study (GWAS) summary statistics for gut microbiota were obtained from the OpenGWAS database, as detailed in Table 2.

**Table 2.**
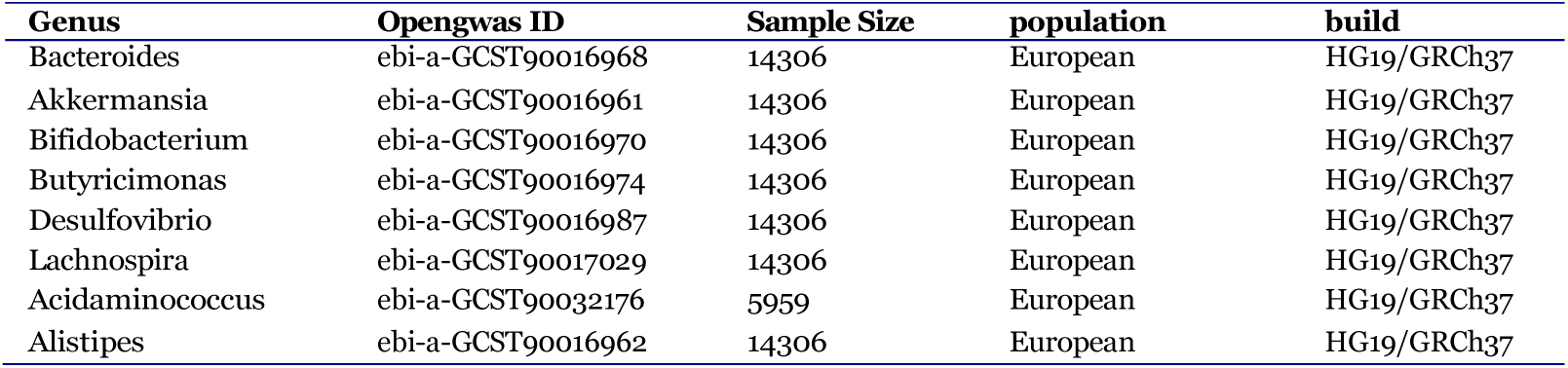
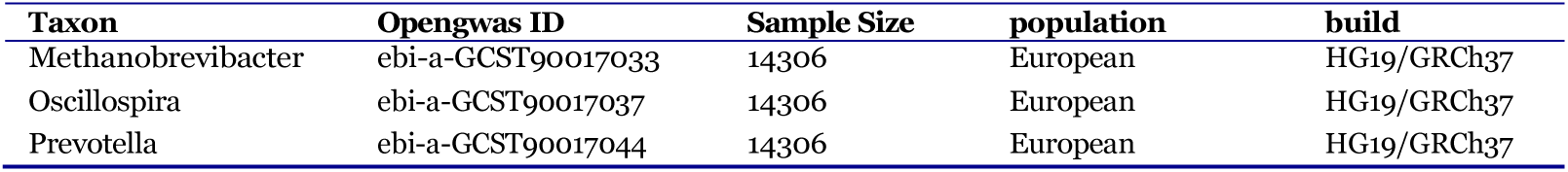
Overview of GWAS datasets for hypertension-associated gut microbiota.

For cohorts without predefined hypertension status, samples were classified into hypertension and healthy control groups according to the World Health Organization (WHO) criteria, defined as systolic blood pressure *≥* 140 mmHg and/or diastolic blood pressure *≥* 90 mmHg. For cohorts with existing phenotype annotations, the original classifications provided in the respective studies were retained.

### Identification of Gut Microbes Robustly Associated with Hypertension

Given the substantial inter-individual variability of the gut microbiota, differential abundance analyses were conducted independently across the three cohorts. Specifically, the Linear Discriminant Analysis Effect Size (LEfSe) method implemented in the microbiomeMarker package[34].was applied to identify differentially abundant microbial genera. Microbial abundance data were normalized using the counts per million (CPM) approach. Statistical significance was defined as a p value < 0.05 combined with an absolute LDA score greater than 2.5. In addition, information on hypertension-associated microbial genera was retrieved from the GutMDisorder database[35]. To enhance the robustness of the findings, the differential genera identified from the three cohorts and the database were intersected. Genera were retained if they were identified in at least two independent cohorts, or if they were detected in one cohort and also reported in the database. Upregulated and downregulated genera were intersected separately to ensure the reliability and consistency of the selected hypertension-associated gut microbes.

### Construction of Gut Microbial Co-occurrence Networks and Topological Analysis

To systematically evaluate the structural roles of hypertension-associated gut microbes within microbial co-occurrence networks, a gut microbial co-occurrence network was constructed based on the CPH16S cohort. First, pairwise correlations among microbial genera were estimated using the SparCC (Sparse Correlations for Compositional data) method[36] to mitigate the compositional effects inherent in microbial abundance data on correlation inference. Subsequently, Random Matrix Theory (RMT) was applied to objectively determine the correlation threshold for network edge inclusion, and an optimal threshold of 0.15 was selected. Based on this threshold, the original correlation network was sparsified to remove weak correlations and reduce noise interference. After obtaining a stable co-occurrence network, the Walktrap community detection algorithm[37] was applied to partition network nodes into modules, thereby identifying network modules enriched with hypertension-associated microbial genera. Finally, node degree centrality was calculated for each genera, and differences in degree between hypertension-associated genera and other microbes were compared. This analysis enabled the assessment of the structural centrality and potential ecological influence of hypertension-associated gut microbes within the co-occurrence network.

### Identification of Genes Involved in Gut Microbiota–Host Interactions and Pathway Enrichment Analysis

To investigate the potential associations between gut microbiota and host genetic background, this study retrieved genome-wide association study (GWAS) summary statistics for nine gut microbial genera significantly associated with hypertension from the OpenGWAS database. Subsequently, MAGMA (Multi-marker Analysis of GenoMic Annotation)[38] was applied to aggregate genera-associated host SNP-level signals to the gene level, yielding gene-based association P values and corresponding Z scores for each gene. To investigate which host biological pathways these microbiota-associated genes primarily participate in, genes were ranked in descending order based on their computed Z-scores, and the top 100 genes most strongly associated with each microbial genera were selected as candidate gene sets to minimize the influence of weakly associated genes. These candidate gene sets were then subjected to Kyoto Encyclopedia of Genes and Genomes (KEGG) pathway enrichment analysis.

### Single-Cell Data Preprocessing and Cell Type Annotation

This study collected peripheral blood mononuclear cell (PBMC) transcriptomic data from 13 control individuals and 12 patients with hypertension[33], comprising a total of 256,403 cells. Cells with a mitochondrial gene expression proportion greater than 10% were excluded. Additional filtering was performed based on the distributions of total UMI counts and the number of detected genes per sample. Doublets were subsequently identified and removed using DoubletFinder, resulting in a final dataset of 199,079 high-quality cells. After quality control, raw UMI count data were log-normalized using the NormalizeData function in Seurat, with a scaling factor of 10,000. Highly variable genes (HVGs) were identified using the variance-stabilizing transformation (VST) method, selecting 2,000 HVGs. Expression values of the selected HVGs were then mean-centered and scaled by their standard deviations using the ScaleData function to minimize the impact of differences in gene expression magnitude on downstream dimensionality reduction analyses. Principal component analysis (PCA) was subsequently performed. To correct for batch effects across samples, the Harmony algorithm was applied in PCA space. The top 30 dimensions were selected for Uniform Manifold Approximation and Projection (UMAP) visualization, construction of the shared nearest-neighbor graph, and cell clustering. Finally, cell types were annotated based on established marker genes (Table S1).

### Quantification of Associations Between Single Cells and Gut Microbiota

To integrate genetic association signals of gut microbiota with single-cell transcriptomic data and systematically characterize potential host–microbe interactions at the cellular level, we applied the scBPS[30] analytical framework to quantitatively assess associations between gut microbial taxa and distinct host cell types. Specifically, based on host–microbe association analyses, gene sets significantly associated with each gut microbial taxon were extracted. Subsequently, a bacterial polygenic score (BPS) was calculated at the single-cell level to quantify the association between each cell and the corresponding microbial taxon. Based on the resulting BPS values for each microbe, all cells were ranked accordingly. Integrating cell type annotations, the area under the curve (AUC) of the BPS ranking curve was calculated for each cell type under a given microbial condition to quantitatively characterize the strength of association between specific gut microbes and host cell types. To assess the statistical significance of these associations, cell orders were randomly permuted 1,000 times, and the BPS-AUC for each cell type was recalculated after each permutation to construct a null distribution, from which empirical significance levels were derived.

### Causal Analysis of Microbial BPS in Relation to Cellular States and Pathway Alterations

Based on the results of the BPS-AUC analysis, the strength and statistical significance of associations between gut microbes and specific cell types can be quantitatively assessed. To further elucidate the potential functional effects of microbes within specific cellular subpopulations, the AUCell method was applied to calculate activity scores of KEGG pathways and Hallmark gene sets at the single-cell level. Subsequently, Pearson correlation analysis was performed at the single-cell resolution to evaluate the relationships between microbial BPS values and corresponding pathway activity scores, thereby revealing microbe-associated changes in cellular functional states. Building upon these analyses, Double Machine Learning (DoubleML)[39] was further employed to investigate the potential causal effects of microbial scBPS on pathway activities within specific cell types. Specifically, microbial scBPS was treated as the treatment variable, while age, sex, and body mass index (BMI) were included as covariates. Pathway activity scores and cellular pseudotime were considered as outcome variables. The DoubleML framework consists of a two-step residualization procedure, in which the conditional expectation functions of the treatment given covariates m(X), and of the outcome given covariates g(X), are estimated separately. In both steps, Random Forest models were used as base learners with the following parameters: num.trees = 100, mtry = 2, min.node.size = 2, and max.depth = 5. After residualization, a linear regression was performed on the residuals of the treatment and outcome variables to estimate the average treatment effect (ATE) and its statistical significance (p value). All resulting p values were adjusted for multiple testing using the false discovery rate (FDR) method.

### Differential Analysis of Cellular Pathway Activity

To investigate differences in pathway activity between host cells from hypertension patients and healthy controls, differential pathway activity analysis was performed using the limma package[40]. Briefly, group variables were encoded using a no-intercept design matrix to separately estimate pathway activity levels in the control and hypertension groups. Linear models were then fitted, and standard errors were moderated using the empirical Bayes approach to improve the stability of differential estimates. Finally, the resulting P values were adjusted for multiple testing using the false discovery rate (FDR) method.

### Group-wise Differential Analysis of Intercellular Communication

Given that the bacterial polygenic score (BPS) of *Butyricimonas* exhibited significant associations across multiple monocyte subtypes, cells were further stratified according to their *Butyricimonas* BPS values and classified into high-BPS and low-BPS groups. On this basis, intercellular communication analyzes were performed separately in each group using the CellChat package[41] to systematically infer ligand–receptor–mediated cell–cell communication networks. Specifically, the number of signaling pathways involved in intercellular communication and the overall interaction strength were calculated for both the high-BPS and low-BPS groups, and corresponding cell–cell communication networks were constructed. Subsequently, differences between the two communication networks were quantitatively assessed to characterize changes in intercellular communication patterns under distinct *Butyricimonas* BPS states.

## Results

### Hypertension-associated gut microbiota display alterations in multiple cohorts and occupy central positions in the network topology

Alpha diversity indices, including Observed, Chao1, ACE, Shannon, Simpson, and Pielou indices, were calculated for each of the three gut microbiome cohorts. The results showed that genera richness–related indices (Observed, Chao1, and ACE) exhibited a decreasing trend in both the CPH16S and CHMG cohorts (Figure S1A–B). In contrast, indices reflecting community diversity and evenness, namely Shannon, Simpson, and Pielou indices, were reduced in the CHMG and CH16S cohorts (Figure S1B–C). Beta diversity analyses revealed statistically significant differences in microbial community structure between hypertension and control groups across all three cohorts (p < 0.05). However, the explained variance for each comparison was relatively low, suggesting that although microbial community compositions were statistically distinguishable, the overall magnitude of the differences was limited (Figure S1D–F). To identify robust microbial biomarkers associated with hypertension, differential abundance analyses at the genus level were performed independently in three cohorts: CPH16S (Figure 1A), CHMG (Figure 1B), and CH16S (Figure 1C), using p < 0.05 and an LDA score > 2.5 as the criteria for significance. To further assess cross-cohort consistency, these results were integrated with hypertension-associated microbial taxa reported in the GutMDisorder database. Genera that were consistently enriched in hypertension (Figure 1D) or consistently depleted in hypertension (Figure 1E) in at least two cohorts, or those that were concordant with database-reported associations, were defined as robust hypertension-associated microbial biomarkers. Using these criteria, *Enterobacter* and *Prevotella* were identified as enriched in hypertension, whereas *Akkermansia, Bifidobacterium, Lachnospira, Acidaminococcus, Butyricimonas, Bacteroides, Oscillospira, Alistipes, Bilophila, Methanobrevibacter, Roseburia*, and *Desulfovibrio* were enriched in healthy controls. Notably, *Methanobrevibacter, Roseburia,* and *Desulfovibrio* did not exhibit significant differences in the CPH16S.

**Figure 1.**
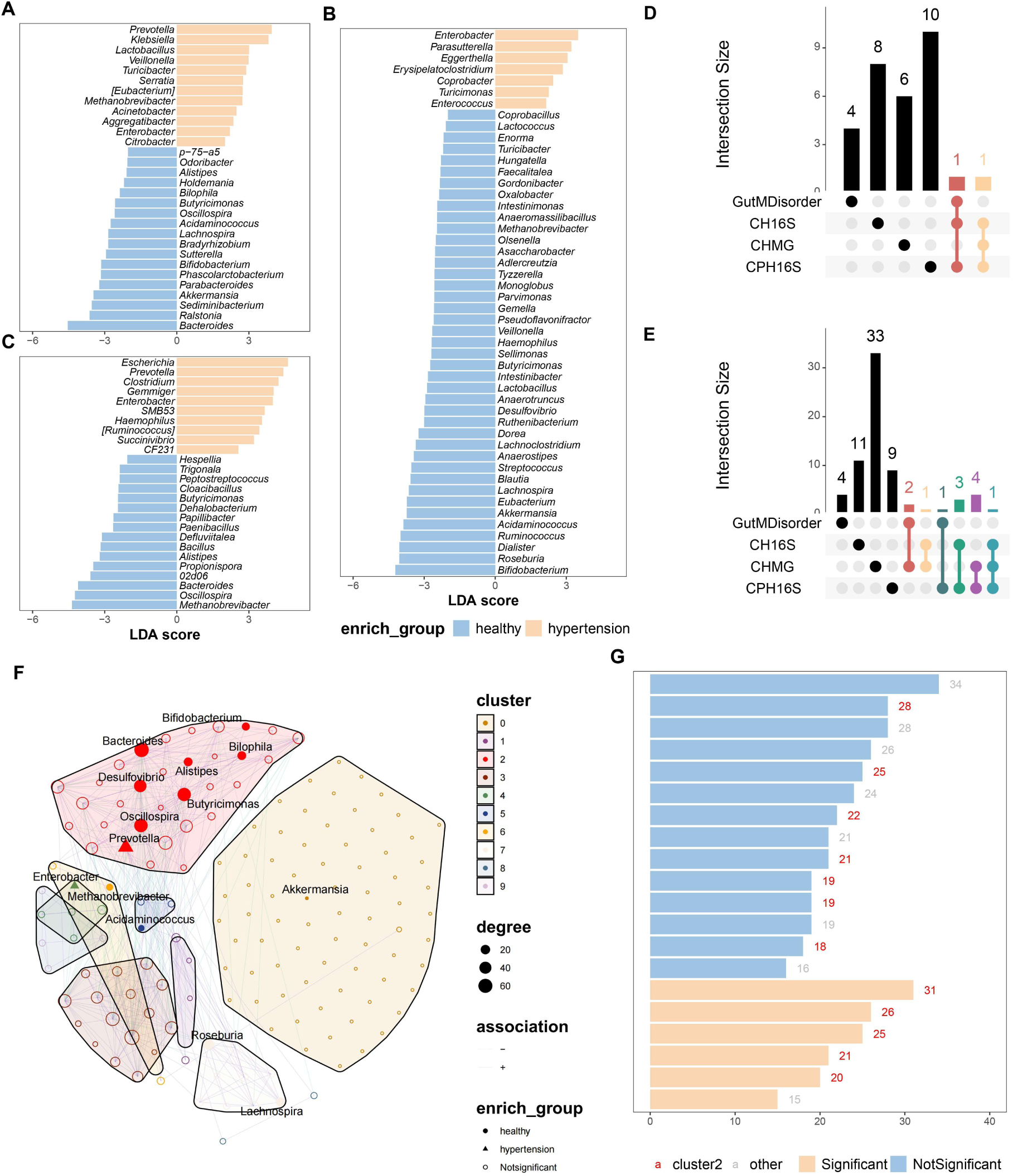
Consistently differential gut microbiota in hypertension and their central positions in the co-occurrence network. Linear Discriminant Analysis (LDA) was used to identify differential microbial taxa between the hypertension and healthy control groups in the CPH16S cohort (A), the CHMG cohort (B), and the CH16S cohort (C). Yellow indicates microbial taxa significantly enriched in the hypertension group, while blue represents taxa enriched in the healthy control group. Subsequently, an intersection analysis was performed on the identified differential taxa across cohorts, resulting in taxa that exhibited consistent enrichment trends in at least two independent cohorts, including those commonly enriched in the hypertension group (D) and those enriched in the healthy control group (E), which were used for further analysis. A co-occurrence network of the microbiota was constructed, followed by network clustering and node connectivity analysis. The size, shape, and color of the nodes represent the node connectivity, enrichment direction, and network cluster affiliation, respectively. The results indicate that the aforementioned differential taxa are predominantly enriched in cluster 2 (F), and the majority of nodes in cluster 2 have connectivity within the top 20 of the entire network (G).

Construction of the gut microbial co-occurrence network revealed that these genera occupied central positions within the network. Community detection further demonstrated that these taxa largely clustered into a single module (Figure 1F). Specifically, 58.3% of the genera enriched in healthy controls were assigned to cluster 2, while 50% of the hypertension-enriched genera also belonged to this cluster (Table S2). Analysis of node degree showed that the degree of this cluster ranked predominantly within the top 20 nodes of the network (Fig. 1G). Furthermore, genera significantly enriched in hypertension were largely associated with other host diseases and exhibited pro-inflammatory potential, whereas taxa capable of producing beneficial metabolites such as short-chain fatty acids and hydrogen sulfide—important for intestinal barrier repair—were depleted in hypertension (Table S3).

### Hypertension-associated robust marker genera are more likely to interact with host immune-related pathways

Based on the previously published study by Kurilshikov A et al., we systematically integrated GWAS summary statistics for the differentially abundant bacterial genera and obtained host genetic association information for nine genera; no usable GWAS data were available for Enterobacter. We then applied the MAGMA framework to integrate genetic associations between each genus and host genes, selecting the top 100 genes ranked by association Z scores for each genus as putative host-associated genes. The results showed that association signals for most genera converged on RBFOX1, PTPRD, and CSMD1 (Figure S2A), suggesting that the genetic regulatory networks involving these genes may indirectly shape gut microbial community structure by modulating intestinal microenvironmental homeostasis. Further KEGG pathway enrichment analysis of the host-associated genes revealed that genes linked to five genera were significantly enriched in the IgSF CAM signaling pathway; genes associated with four genera were significantly enriched in the Rap1 signaling pathway, Cell adhesion molecule (CAM) interactions, and Arrhythmogenic right ventricular cardiomyopathy; and genes associated with three genera were significantly enriched in pathways such as Bacterial invasion of epithelial cells and Human papillomavirus infection (Figure S2B).

### Interactions between gut microbiota and peripheral blood cells modulate host blood pressure

To investigate the interactions between gut microbiota and host peripheral blood mononuclear cells (PBMCs), we collected PBMC transcriptomic data from 13 control samples and 12 patients with hypertension, comprising a total of 256,403 cells. After doublet removal and filtering of low-quality cells, 199,079 high-quality cells were retained for downstream analyses (Figure S3A–B). The data was normalized and corrected for batch effects (Figure S3C–D). Cells were then classified by clustering analysis (resolution = 0.8), resulting in the identification of 37 cell clusters (Figure 2A). Based on cell-type markers curated in the CellMarker database, these clusters were annotated into 12 distinct cell types, including progenitor neutrophils (ProNeutrophils), hematopoietic stem cells (HSCs), plasma cells (PlasmaCells), B cells (B cells), T cells (T cells), natural killer (NK) cells, neutrophils, basophils, monocytes, plasmacytoid dendritic cells (pDCs), erythrocytes, and platelets (Figure 2B–C). Among these cell types, NK cells and pDCs were significantly reduced in proportion in the hypertension group, whereas platelets were significantly increased (Figure S4A). Further differential expression analysis revealed that NK cells exhibited the largest number of differentially expressed genes, which were significantly enriched in multiple immune- and inflammation-related pathways, including IL-17 signaling pathway, Th1 and Th2 cell differentiation, and Cell adhesion molecule (CAM) interactions, as well as several cardiovascular-related pathways such as Dilated cardiomyopathy and Type I diabetes mellitus (Figure S4B, E). In contrast, pDCs displayed relatively few transcriptional differences between groups, with their differentially expressed genes mainly enriched in the Coronavirus disease-COVID-19 pathway (Figure S4C, F). There are some differentially expressed genes in platelets, but they are not enriched in KEGG pathways (Figure S4D).

**Figure 2.**
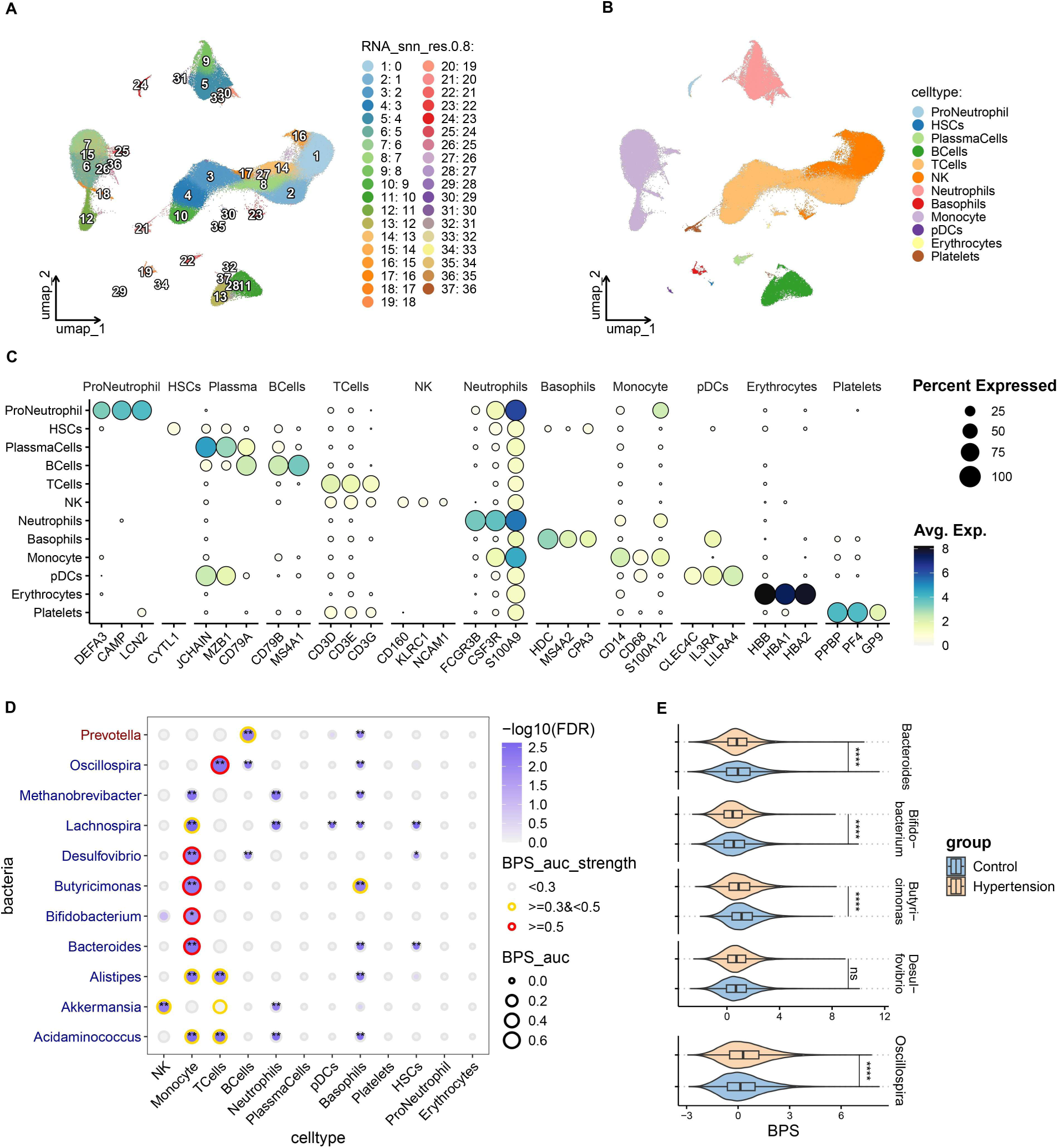
Interactions between hypertension-associated gut microbiota and peripheral blood mononuclear cells. Unsupervised clustering analysis of peripheral blood cells based on single-cell transcriptomic data, with different colors representing distinct cell clusters (A). Cell type annotation of each cluster based on known cell-specific marker genes, with different colors indicating different cell types (B). Visualization of the expression distribution of representative cell marker genes (C). The scBPS method was then used to assess the association between robustly differentially abundant microbiota and various cell types, with bubble size and color representing the strength and statistical significance of the association, respectively. The outer color of the bubble indicates the strength level (red ≥ 0.5, gray < 0.3, yellow for intermediate range) (D). Finally, a group comparison analysis of the top 5 microbiota with the strongest association to each cell type was conducted, where the x-axis represents the microbiota-cell association strength (BPS value) and the y-axis represents different cell cluster statuses (E).

Using single-cell bacterial polygenic score (scBPS) analysis, we identified varying degrees of association between specific bacterial genera and particular cell types, including Prevotella, Bacteroides, Akkermansia, Bifidobacterium, Butyricimonas, Oscillospira, Alistipes, Methanobrevibacter, and Desulfovibrio. Notably, Bacteroides, Bifidobacterium, Butyricimonas, and Desulfovibrio showed BPS scores greater than 0.5 in monocytes, whereas Oscillospira exhibited a BPS score greater than 0.5 in T cells (Figure 2D). Furthermore, the BPS scores of Bacteroides, Bifidobacterium, and Butyricimonas in monocytes were significantly lower in the hypertension group than in controls, while the BPS score of Oscillospira in T cells was significantly higher in the hypertension group. Notably, no significant difference in the abundance of Desulfovibrio was observed in the CPH16S cohort, and its BPS score in monocytes did not differ significantly between the hypertension and control groups (Figure 2E).

### *Bacteroides* and *Bifidobacterium* associated with proliferation- and repair-related functions in classical monocytes

Monocytes were subjected to re-clustering analysis, resulting in the identification of 18 distinct cell clusters (Figure 3A). These clusters were re-annotated according to established marker genes for classical, intermediate, and non-classical monocytes. Cluster 7 was characterized by the absence of FCGR3A and CD14 expression, together with high expression levels of CST3 and S100A8 (Figure 3B–C), and was therefore defined as a non-classical–like monocyte population. Ultimately, we annotated 30,961 classical monocytes, 2,951 intermediate monocytes, 5,886 non-classical monocytes, and 1,358 non-classical–like monocytes (Figure 3D). Across these monocyte subpopulations, the bacterial polygenic scores (BPS) of *Bacteroides, Bifidobacterium, Butyricimonas*, and *Desulfovibrio* reached their highest levels in classical monocytes (Figure 3E). Moreover, the BPSs of *Bacteroides*, *Bifidobacterium*, and *Butyricimonas* in classical monocytes were significantly lower in patients with hypertension than in controls. Similarly, the BPS of *Butyricimonas* was also significantly reduced in other monocyte subtypes in the hypertension group compared with controls (Figure 3F). To further elucidate the functional relevance of these associations, AUCell was used to calculate KEGG pathway activity scores in classical monocytes, and the correlations between pathway activities and the BPSs of *Bacteroides* and *Bifidobacterium* were examined. The results revealed a high degree of similarity between the BPSs of *Bacteroides* and *Bifidobacterium*. In particular,kegg beta alanine metabolism, kegg drug metabolism other enzymes, and kegg pantothenate and COA biosynthesis showed the strongest correlations with the BPSs of both genera (Figure 3G). Consistent results were obtained when pathway scores from other databases were analyzed: the BPSs of *Bacteroides* (Figure 3H) and *Bifidobacterium* (Figure 3I) were significantly positively correlated with pyrimidine/nucleotide metabolism–related pathways, including reactome pyrimidine catabolism, wp pyrimidine metabolism and related diseases, wp biomarkers for pyrimidine metabolism disorders, wp fluoropyrimidine activity and reactome nucleotide catabolism. In addition, the BPSs of *Bacteroides* and *Bifidobacterium* were also significantly positively correlated with EGF-EGFR signaling. In addition, protein–protein interaction (PPI) network analysis revealed that host genes associated with *Bacteroides* and *Bifidobacterium*, identified through MAGMA integration, could regulate genes involved in the EGFR signaling pathway (Figure S5A,B). These findings suggest that Bacteroides and *Bifidobacterium* may regulate processes such as proliferation, repair, and activation of classical monocytes.

**Figure 3.**
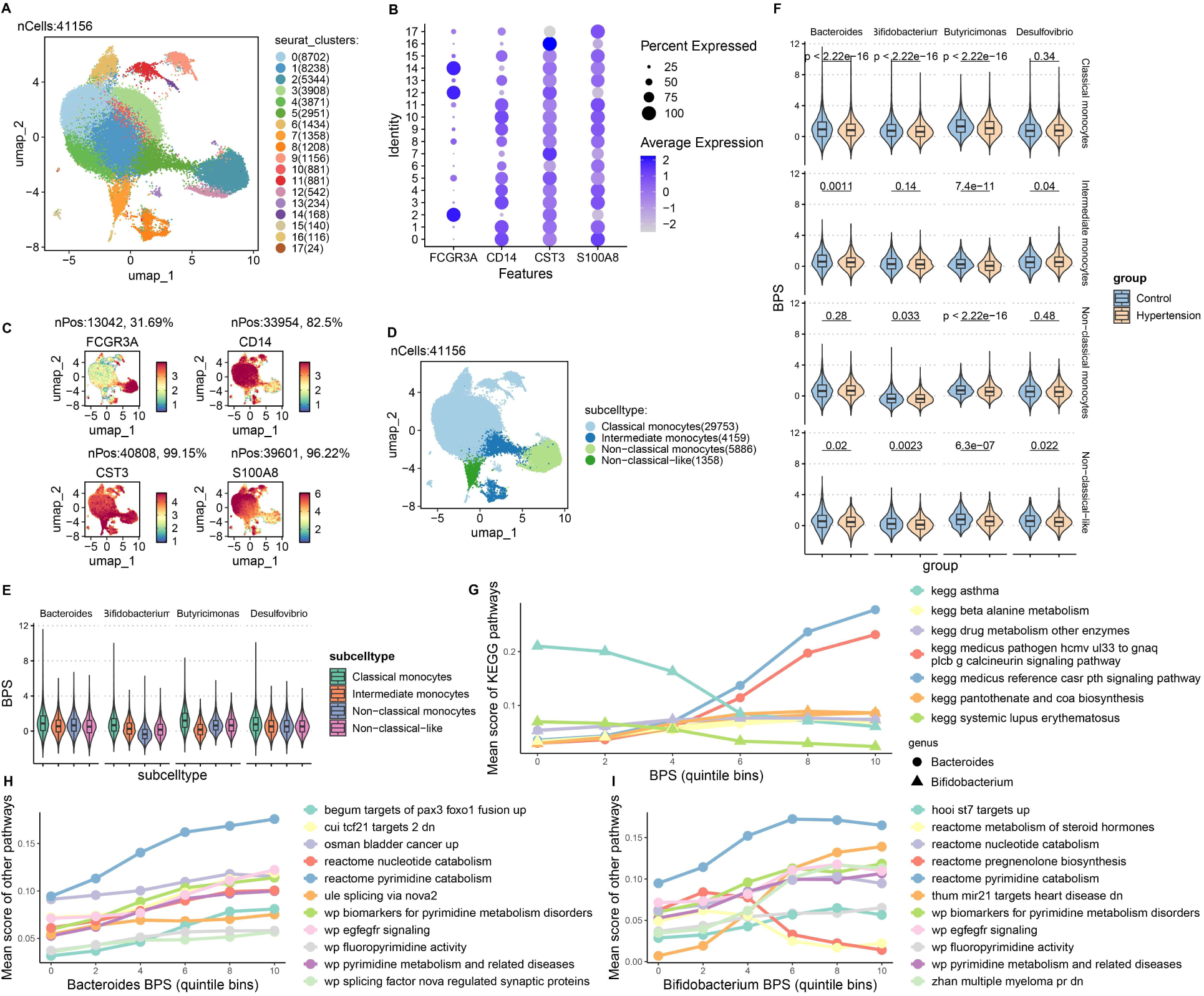
Effects of *Bacteroides* and *Bifidobacterium* on the functional states of monocyte. UMAP projection of re-clustered monocyte subgroups, with different colors representing distinct subgroups (A). The average expression levels and expression frequencies of monocyte subtype marker genes (B). Visualization of the expression distribution of the above marker genes in the UMAP space (C). Overall distribution and cell count statistics of monocyte subgroups in the UMAP space (D). The distribution of the BPS (microbiota-cell association) between Bacteroides, Bifidobacterium, Butyricimonas, and Desulfovibrio across different monocyte subgroups. The y-axis represents the genera-cell BPS, and the color indicates different monocyte subtypes (E). Differential analysis of microbiota BPS across different monocyte subtypes (F). Correlation analysis between monocyte KEGG pathway activity scores and BPS of Bacteroides and Bifidobacterium. The top 5 pathways with the highest correlation are shown, with circles and triangles representing Bacteroides and Bifidobacterium, respectively, and the color representing different pathways (G). Further pathway activity scores were calculated based on other functional databases and their correlation with BPS of Bacteroides (H) and Bifidobacterium (I), displaying the top 10 pathways with the highest correlation.

### *Butyricimonas* is involved in regulating antigen presentation in host classical monocytes

Correlation analyses were performed between the bacterial polygenic score (BPS) of *Butyricimonas* and KEGG pathway activities in classical, intermediate, and non-classical monocytes (Figure 4A–C). In classical monocytes, *Butyricimonas* BPS showed significant negative correlations with multiple antigen presentation and immune-related pathways, including antigen processing and presentation, medicus reference antigen processing and presentation by MHC class II molecules and intestinal immune network for IGA production (Figure 4A). In intermediate monocytes, medicus reference antigen processing and presentation by MHC class II molecules also exhibited a significant negative correlation with *Butyricimonas* BPS. In addition, several pathways related to mitochondrial energy metabolism and oxidative stress (such as parkinsons disease adn huntingtons disease), as well as the graft versus host disease pathway reflecting T cell-mediated immune activation, were significantly negatively correlated with *Butyricimonas* BPS (Figure 4B). In non-classical monocytes, *Butyricimonas* BPS was likewise mainly negatively correlated with pathways associated with mitochondrial energy metabolism and oxidative stress, including parkinsons disease and huntingtons disease (Figure 4C). PPI network analysis revealed that host genes associated with *Butyricimonas*, identified through MAGMA integration, could regulate genes involved in the antigen processing and presentation pathway (Figure S5C). Given that antigen presentation is predominantly mediated by intermediate monocytes, pseudotime analysis was further performed to investigate the dynamic transitions among monocyte subpopulations. The results indicated that classical monocytes progressively differentiate into intermediate monocytes along the pseudotime trajectory (Figure 4D–E). During this transition, the activity of antigen presentation–related pathways showed a marked increasing trend, and throughout the entire trajectory, pathway activity was consistently higher in cells from the hypertension group than in those from the control group (Figure 4F–G). Furthermore, causal inference analysis was conducted using the DoubleML framework, with *Butyricimonas* BPS as the treatment variable and antigen presentation pathway activity and pseudotime progression as outcome variables. The results revealed significant negative average treatment effects (ATE) of *Butyricimonas* BPS on both outcomes, suggesting that *Butyricimonas*-associated signals may exert an inhibitory effect on monocyte antigen presentation functions and on the differentiation of classical monocytes toward the intermediate state (Figure 4H).

**Figure 4.**
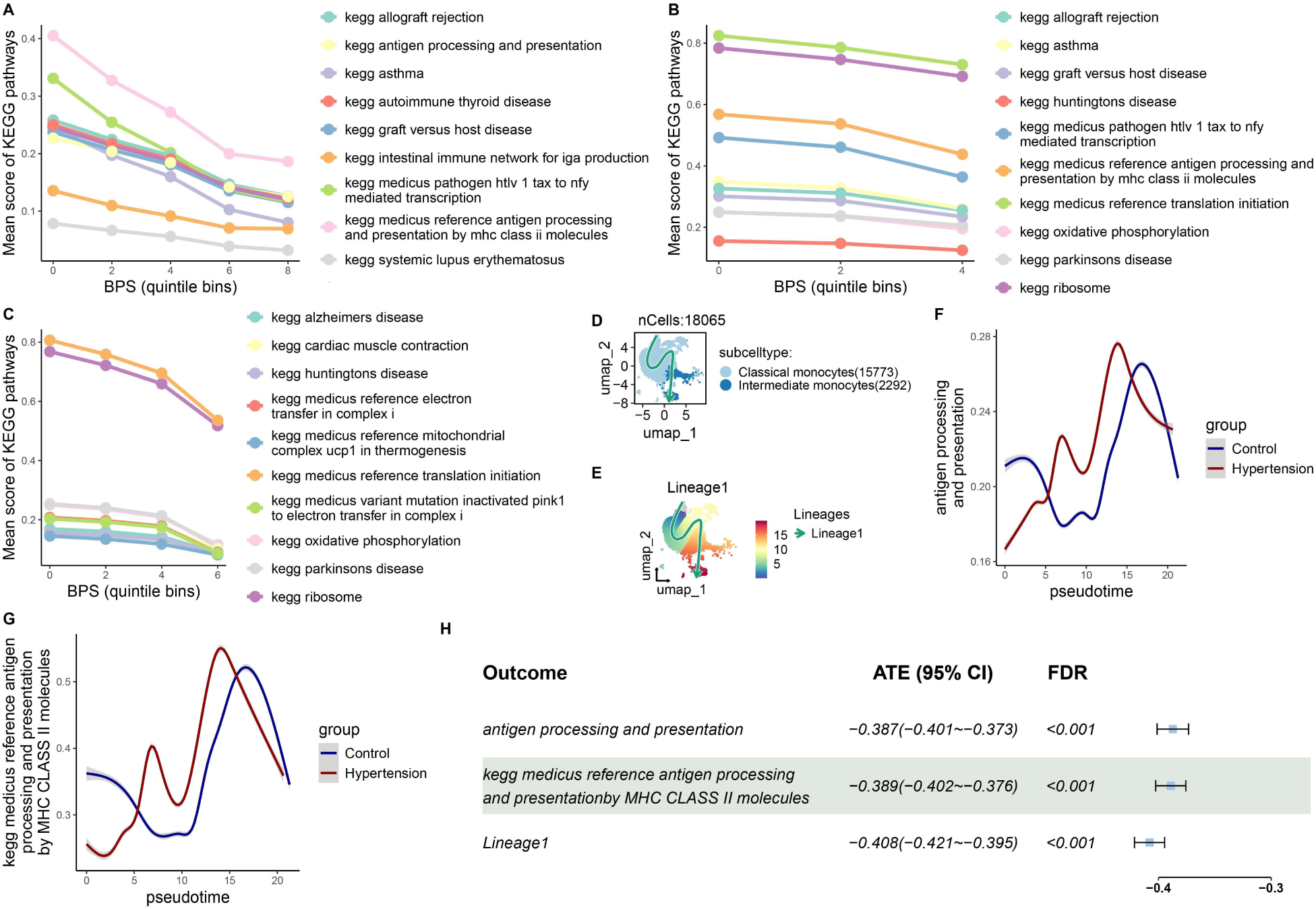
Effect of *Butyricimonas* on the functional state of monocytes. KEGG pathway activity scores and their correlation with *Butyricimonas* BPS in classical monocytes (A), intermediate monocytes (B), and non-classical monocytes (C), where colors indicate variations in different pathways. Differentiation trajectories of classical and intermediate monocytes in the hypertension group (D-E). Dynamic changes in antigen presentation-related pathway activity along pseudo-time trajectories, with red representing the hypertension group and blue representing healthy controls (F-G). Causal relationship between *Butyricimonas* BPS, cell trajectories, and antigen presentation pathway activity inferred using DoubleML, with ATE representing the average treatment effect of *Butyricimonas* BPS on cell trajectories and antigen presentation pathway activity (H).

### *Butyricimonas* is associated with intercellular communication among host immune cells

Cell–cell communication analysis of samples from the hypertension and control groups revealed that, compared with controls, the global communication network in the hypertension group was markedly more active, as reflected by significant increases in both the number of signaling pathways and the overall interaction strength. Specifically, a total of 2,895 signaling interactions were detected in the hypertension group, with an overall interaction strength of 119.656, whereas only 2,290 signaling interactions with an interaction strength of 93.046 were identified in the control group (Figure S6A). The hypertension samples were further stratified into High BPS and Low BPS groups based on *Butyricimonas* BPS levels. The High BPS group exhibited 2,406 signaling interactions with an interaction strength of 108.08, whereas the Low BPS group showed 3,111 signaling interactions and a markedly higher interaction strength of 137.473. These results indicate that, in the context of hypertension, lower *Butyricimonas* BPS is associated with more frequent and stronger intercellular communication (Figure 5A). Differential analysis of the communication networks between the hypertension and control groups showed that nearly all cell–cell pairs in the hypertension group displayed increased numbers of interactions, accompanied by a global enhancement of interaction strength (Figure S6B–C). Consistently, within the hypertension group, most cell–cell pairs in the High BPS group exhibited fewer interactions and lower interaction strength than those in the Low BPS group (Figure 5B–C). Notably, MHC-II–related signaling pathways targeting CD4 T cells emerged exclusively in the hypertension group and were not detected in the control group (Figure S6D). Further stratified analysis revealed that within the hypertension group, only cells in the Low BPS group displayed MHC-II–mediated signaling toward CD4 T cells, whereas such interactions were absent in the High BPS group (Figure 5D). These findings suggest that *Butyricimonas* BPS levels may be involved in regulating antigen presentation–related immune cell communication patterns.

**Figure 5.**
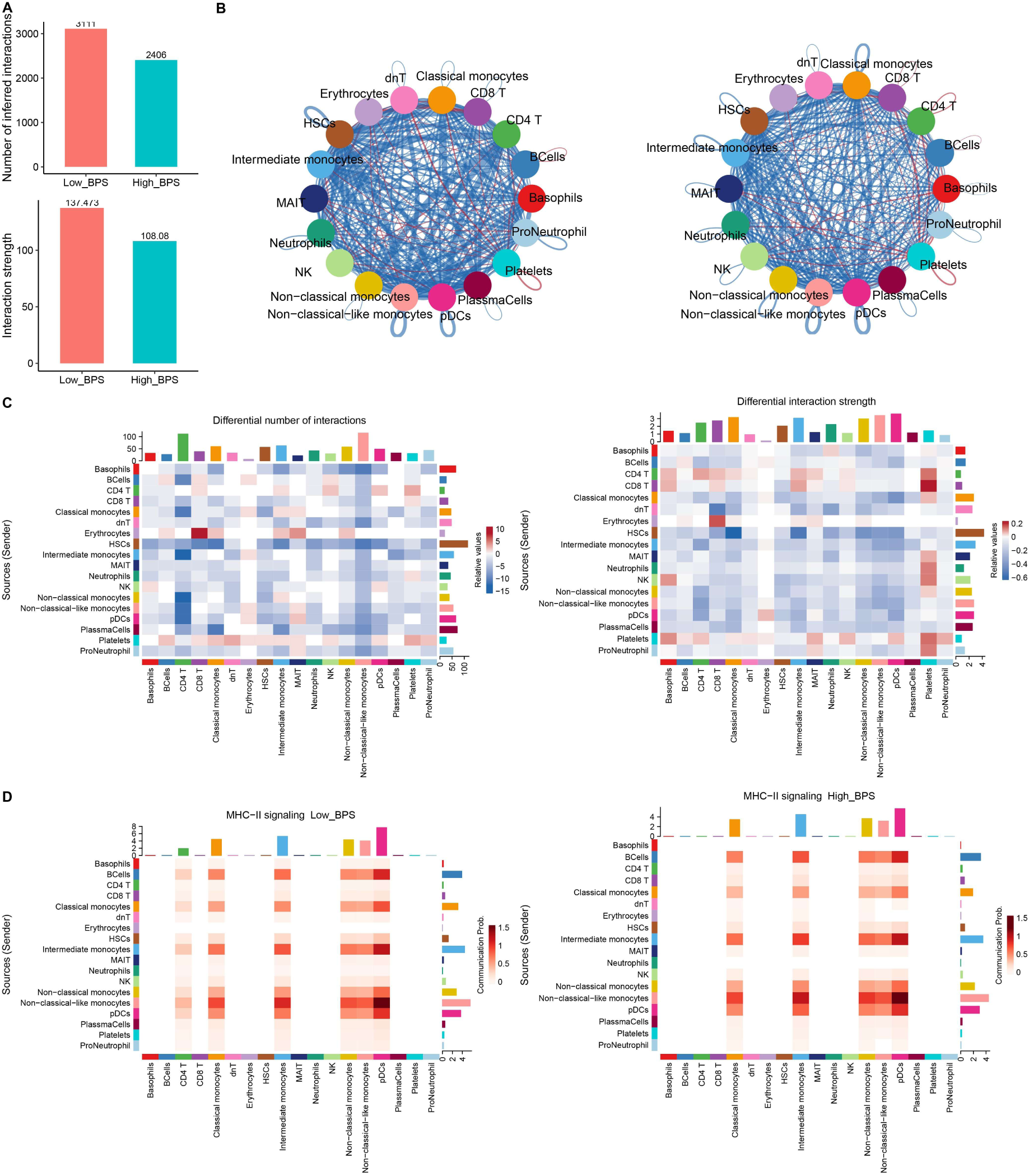
Effect of *Butyricimonas* on intercellular communication among immune cells. Overall comparison of cell communication quantity and intensity between the high and low *Butyricimonas* BPS groups (A). Network plot of the differences in cell communication quantity and intensity, where nodes represent different immune cell types, and edges indicate intercellular communication relationships (B). Heatmap of the differences in cell communication quantity and intensity, with red indicating significantly upregulated communication in the high *Butyricimonas* BPS group and blue indicating significantly downregulated communication in the high *Butyricimonas* BPS group (C). Differential cell communication mediated by the MHC-II signaling pathway between the high and low *Butyricimonas* BPS groups (D).

### *Oscillospira* is associated with dnT cell–mediated immunosuppression and inflammatory responses

Re-clustering analysis of T cells identified 18 distinct cell clusters (Figure 6A), which were subsequently annotated based on canonical marker genes of known T-cell subpopulations (Figure 6C). In total, 11 T-cell subsets were identified, including 16,170 CD4 Naive, 7,632 CD4 TCM, 4,695 CD4 TEM, 3,898 CD4 CTL, 1,732 Treg, 2,269 MAIT, 8,365 CD8 Naive, 5,442 CD8 TCM, 7,075 CD8 TEM, 16,463 CD8 CTL, and 1,043 dnT cells (Figure 6B). Comparative analysis of cell proportions revealed that, relative to controls, the hypertension group showed a significant decrease in CD8 Naive and dnT cell proportions, whereas CD8 TCM cells were significantly increased (Figure 6D). Analysis of the distribution of *Oscillospira* bacterial polygenic scores (BPS) across T-cell subsets revealed that *Oscillospira* BPS was highest in dnT cells, followed by relatively high levels in CD4 Naive, CD8 Naive, CD4 TCM, and CD8 TCM cells (Figure 6E). Notably, dnT cells in the hypertension group exhibited significantly higher *Oscillospira* BPS compared with controls (Figure 6F). Using the AUCell method, Hallmark pathway activities were calculated for dnT cells and correlated with *Oscillospira* BPS. The results showed a significant negative correlation between *Oscillospira* BPS and the inflammatory response pathway, but a significant positive correlation with the IL2 STAT5 signaling pathway in dnT cells (Figure 6G). PPI network analysis revealed that host genes associated with *Oscillospira*, identified through MAGMA integration, could regulate genes involved in the IL2 STAT5 signaling and inflammatory response pathways (Figure S5D,E). Further comparison of pathway activities between groups revealed that the inflammatory response pathway was significantly downregulated in dnT cells from the hypertension group (Figure 6H), whereas in other T-cell subsets—including CD4 Naive, CD8 Naive, CD4 TCM, and CD8 TCM cells—this pathway was generally upregulated (Figure 6I). These findings suggest that dnT cells exhibit distinct functional characteristics under hypertensive conditions, shifting toward an immunoregulatory phenotype rather than a pro-inflammatory state. Based on IL2 STAT5 signaling activity, dnT cells were divided into IL2 STAT5-high and IL2 STAT5-low subpopulations, and pseudotime analysis revealed a differentiation trajectory from low to high IL2 STAT5 states (Figure 6J–K). Consistent with this, the overall proportion of dnT cells was markedly reduced in the hypertension group (Figure 6D).

**Figure 6.**
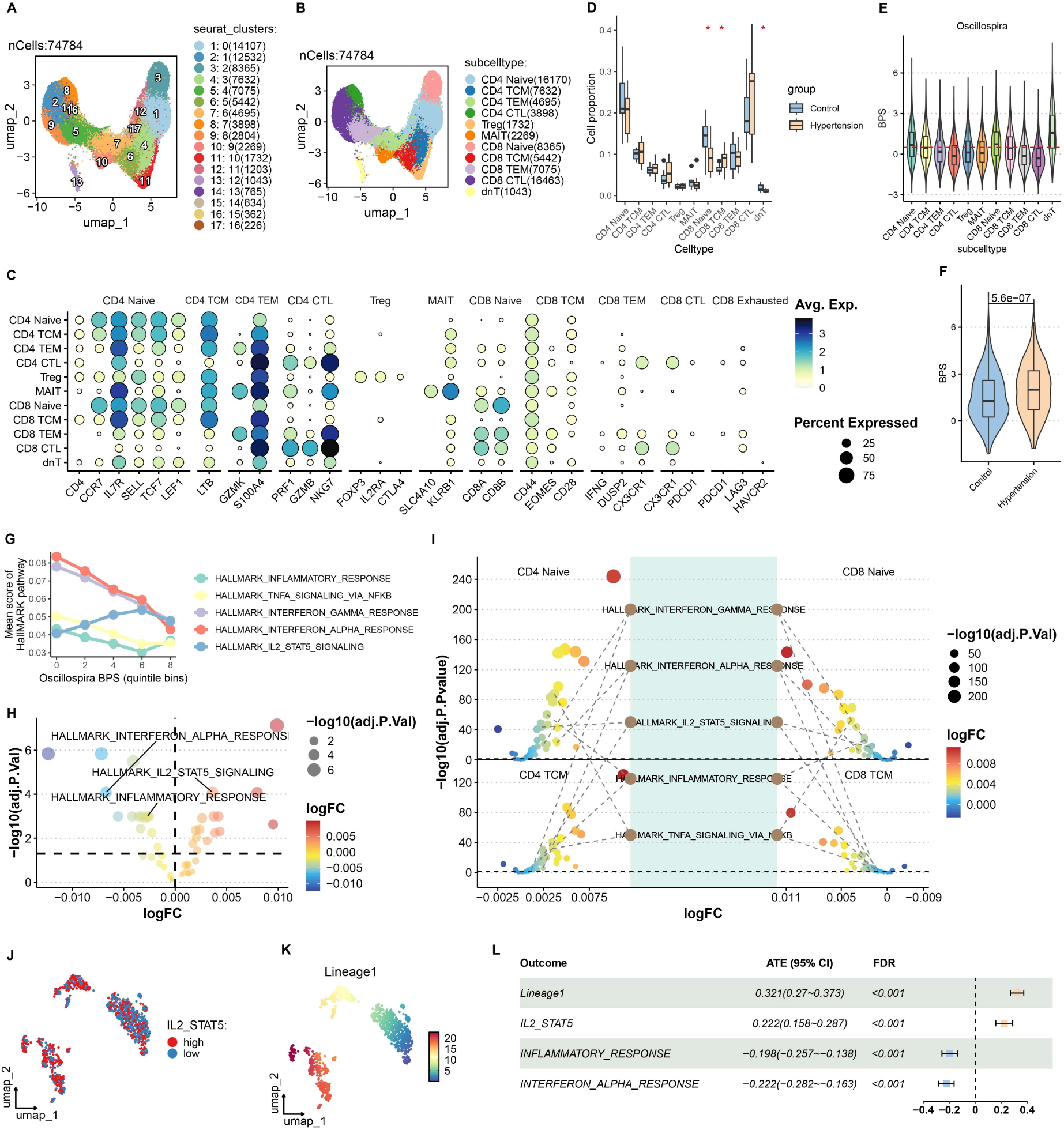
Effect of *Oscillospira* on the functional state of dnT cells. Unsupervised re-clustering of T cells, with UMAP dimensionality reduction showing different T cell subpopulations, where colors represent distinct cell clusters (A). Cell type annotation of re-clustered cells based on T cell subtype markers, with UMAP projection, where different colors represent various T cell subtypes (B). Bubble plot showing the average expression levels and the proportion of cells expressing each T cell subtype marker (C). Comparison of the composition ratio differences of different T cell subtypes between the hypertension and healthy control groups, with yellow indicating the hypertension group, blue indicating the healthy control group, and asterisks indicating statistically significant differences (*p* < 0.05 ∗) (D). Distribution of Oscillospira BPS across different T cell subtypes(E), and its differential expression in dnT cells between the hypertension and healthy control groups (F). Correlation analysis between the Hallmark pathway activity scores in dnT cells and Oscillospira BPS, with colors representing different functional pathways (G). Differential gene expression analysis in dnT cells, where bubble color and size represent significance level and *log*2 fold change, respectively (H). Differential gene expression analysis in CD4 Naive, CD8 Naive, CD4 TCM, and CD8 TCM cells (I). Dimensionality reduction analysis of dnT cells, with classification into high and low IL2-STAT5 pathway activity groups based on IL2-STAT5 pathway activity (J). Further inference of the differentiation trajectory of the two dnT cell groups, with colors representing pseudo-time progress (K). Finally, causal inference of Oscillospira BPS on the dnT cell pseudo-time trajectory, IL2-STAT5 pathway activity, and inflammation-related pathway activities using the DoubleML method (L).

Finally, causal inference analysis was performed using the DoubleML framework, with *Oscillospira* BPS as the treatment variable and dnT pseudotime progression, IL2 STAT5 signaling, and inflammation-related pathways as outcome variables. The results showed that *Oscillospira* BPS had a significant positive ATE on both dnT pseudotime progression and IL2 STAT5 signaling activity, whereas it exhibited significant negative ATEs on inflammatory response and interffron alpha response pathways (Figure 6L). These findings indicate that *Oscillospira*-associated signals may promote an IL2 STAT5–driven immunosuppressive phenotype in dnT cells while suppressing inflammatory activation under hypertensive conditions.

## Discussion

This study systematically dissected gut microbiota dysbiosis and its interactions with the host immune system under hypertensive conditions across multiple analytical layers, including community structure, ecological network topology, host–microbiome genetic associations, single-cell functional profiling, and intercellular communication. Our findings further demonstrate that hypertension is accompanied by a functional reorganization of the gut microbial ecosystem that is tightly coupled with dysregulated host immune regulation. Together, these results provide a more refined and dynamic framework for understanding the mechanistic roles of the gut microbiota in the initiation and progression of hypertension. Differential abundance analysis identified multiple genera with altered abundance in the hypertensive group. Among these, *Enterobacter* and *Prevotella* have been implicated in various diseases and exhibit pro-inflammatory potential, whereas several beneficial genera—such as *Akkermansia, Bifidobacterium, Lachnospira, Acidaminococcus*, and *Butyricimonas*—which are capable of producing short-chain fatty acids and hydrogen sulfide that support intestinal barrier repair, were depleted or reduced in abundance in the hypertensive gut. This pattern suggests that hypertension may be associated with the loss of potentially protective or regulatory microbial taxa, thereby impairing the functional capacity of the gut microbiota to maintain host blood pressure homeostasis, consistent with previous reports[42, 43, 44]. Moreover, gut microbial co-occurrence network analysis revealed that these taxa exhibited higher network degree, indicating that they occupy relatively central topological positions within the gut microbial ecosystem. Network nodes degree reflects the number and strength of potential interactions between a given genera and other microbial members; taxa with higher degree are generally more engaged in direct or indirect ecological interactions and are therefore likely to play critical roles in maintaining community stability and functional coordination[45, 46]. To further investigate the potential interactions between the identified key microbial genera and the host transcriptome, we applied the MAGMA framework to integrate microbial features with host genetic association signals. The results revealed that these microbial genera exhibited significant associations across multiple host genetic loci, with signals predominantly enriched in genomic regions harboring RBFOX1, PTPRD, and CSMD1. Among these, CSMD1 has been reported to be involved in inflammatory responses[47]. while PTPRD, a member of the protein tyrosine phosphatase family, functions as a critical signaling molecule regulating diverse cellular processes, including cell growth and differentiation. These findings suggest that microbiota-associated signals may influence the functional state of peripheral immune cells through host genetic susceptibility. In microbiota–host cell interaction analyses, we observed significant associations between *Bacteroides, Bifidobacterium, Butyricimonas, Oscillospira*, and *Desulfovibrio* with peripheral blood monocytes and T cells, indicating that these gut microbes may participate in peripheral immune regulation through genetic or molecular interactions. Previous studies have shown that these genera are commonly involved in short-chain fatty acid production, bile acid metabolism, or inflammatory signaling regulation[48, 49, 50, 51]. Their metabolic products, such as butyrate, can modulate monocyte inflammatory responses and influence T cell differentiation and functional states[52, 53]. thereby contributing to the maintenance of host immune homeostasis. Furthermore, several genera exhibited significant differences in bacterial–phenotype scores (BPS) between hypertensive and control groups, suggesting that these microbes are not only genetically associated with host immune cells but may also undergo disease-related alterations in interaction strength or functional states. This observation further supports previous findings that gut microbiota dysbiosis can indirectly contribute to hypertension and cardiovascular disease progression by activating immune responses and sustaining chronic low-grade inflammation[54]. However, although *Desulfovibrio* displayed genetic association signals with monocytes and T cells, no significant abundance differences were observed between hypertensive and control individuals in the CPH16S prospective cohort, nor did its BPS differ significantly between groups. This suggests that *Desulfovibrio* may represent a genetically associated microbial taxon rather than a key functional genera directly involved in host gene regulation or immune interaction dynamics under the conditions of this study. Importantly, distinct gut microbes exhibited heterogeneous interaction patterns at the immune cell level. *Butyricimonas*, in particular, demonstrated a more specific and mechanistically directed immune interaction profile. Our analysis revealed a significant negative association between *Butyricimonas* and antigen processing and presentation pathways in classical and intermediate monocytes. This finding suggests that *Butyricimonas* may promote intestinal barrier repair and enhance epithelial integrity through the secretion of short-chain fatty acids such as butyrate[50], thereby reducing systemic exposure to gut-derived antigens. Consequently, this may attenuate activation of monocyte antigen presentation pathways and limit the transmission of immune activation signals to downstream immune cells, such as T cells[55, 56]. Consistently, cell–cell communication analysis revealed weaker antigen presentation signaling in cells with higher *Butyricimonas* BPS. Together, these results suggest that *Butyricimonas* may exert an inhibitory regulatory role during the transition from innate to adaptive immunity, helping to constrain excessive immune activation and prevent the establishment of chronic inflammatory states.

Additionally, *Oscillospira* was found to promote the differentiation of dnT cells toward regulatory immune phenotypes, suggesting its potential involvement in modulating immune tolerance and immunosuppressive balance within the peripheral immune system. Given that *Oscillospira* has frequently been associated with metabolic health and anti-inflammatory states in previous studies[57, 58], this finding further supports the notion that Oscillospira may indirectly shape systemic immune homeostasis by influencing immune cell differentiation trajectories.

By integrating gut microbiome profiling, robust multi-cohort screening, host genetic association analyses, and single-cell transcriptomic data, this study systematically elucidates the multi-layered mechanisms underlying gut microbiota dysbiosis and its interactions with host immune cells in the context of hypertension. Our findings demonstrate that hypertension-associated microbial taxa occupy key topological positions within ecological networks and may influence the functional states of peripheral immune cells through host genetic pathways, particularly in processes related to monocyte antigen presentation and T cell–mediated immune regulation. By incorporating a single-cell bacterial–phenotype score (scBPS) framework and causal inference approaches, this study characterizes the functional directionality of microbiota–host immune interactions at single-cell resolution. These results provide novel immunological evidence supporting the role of gut microbiota in the initiation and progression of hypertension. Collectively, our findings not only extend the theoretical framework of microbiome research in hypertension but also identify potential targets and a conceptual basis for precision intervention strategies centered on specific microbiota–immune axes.

It should be noted, however, that although this study employed the doubleML method, which is designed for causal inference, to explore potential causal relationships between microbiota and cellular functions, the ATE computed by this method cannot be directly interpreted as the true causal effect. Rather, it primarily reflects the estimated effect trend under the assumptions of the model and the available data, rather than a definitive causal conclusion.

Furthermore, several other limitations should be noted. First, this study is primarily based on cross-sectional data, which limits the ability to infer temporal relationships and causal directions among gut microbiota alterations, immune dysregulation, and blood pressure elevation. Second, although most of the microbiome and single-cell transcriptomic data were obtained from Chinese participants, the lack of GWAS summary data for Asian populations required the use of European GWAS summary data, which may introduce a certain degree of bias. Third, due to the lack of single-cell transcriptomic data from gut epithelial or mucosal immune cells under hypertensive conditions, our analyses mainly relied on peripheral blood immune cells, and thus primarily reflect systemic rather than local intestinal immune interactions. Finally, the use of 16S rRNA sequencing restricts taxonomic resolution, precluding the precise characterization of species- or strain-level functional heterogeneity.

Future studies integrating longitudinal designs, functional intervention experiments, and higher-resolution multi-omics approaches—such as shotgun metagenomics, metatranscriptomics, and intestinal epithelial or spatial transcriptomics—will be essential to validate the causal roles of key microbial taxa and immune pathways. Such efforts will further strengthen the mechanistic understanding of micro-biota–immune interactions in hypertension and facilitate the development of microbiome-based precision intervention strategies.

## Conclusion

This study systematically elucidates the potential mechanisms underlying gut microbiota dysbiosis and its interactions with the host immune system in the context of hypertension by integrating gut microbiome profiling, multi-cohort robustness analyses, host genetic association analyses, single-cell transcriptomics, and cell–cell communication data. The results indicate that hypertension is characterized predominantly by a reconstruction of microbial ecological functions and the loss of key functional taxa, which occupy central topological positions within ecological networks and may influence the activation, adhesion, antigen presentation, and immunoregulatory processes of peripheral immune cells through host genetic pathways. Furthermore, at single-cell resolution, this study reveals directional associations between specific microbes (such as *Butyricimonas* and *Oscillospira*) and the functional states of monocytes and T cells, suggesting that gut microbiota may contribute to the onset and progression of hypertension by modulating immune homeostasis (Figure 7). Collectively, these findings extend the theoretical framework of hypertension-related microbiome research from compositional alterations to functional and immune interaction mechanisms, and provide an important theoretical basis and potential research directions for precision interventions and personalized prevention strategies targeting specific microbiota–immune axes.

**Figure 7.**
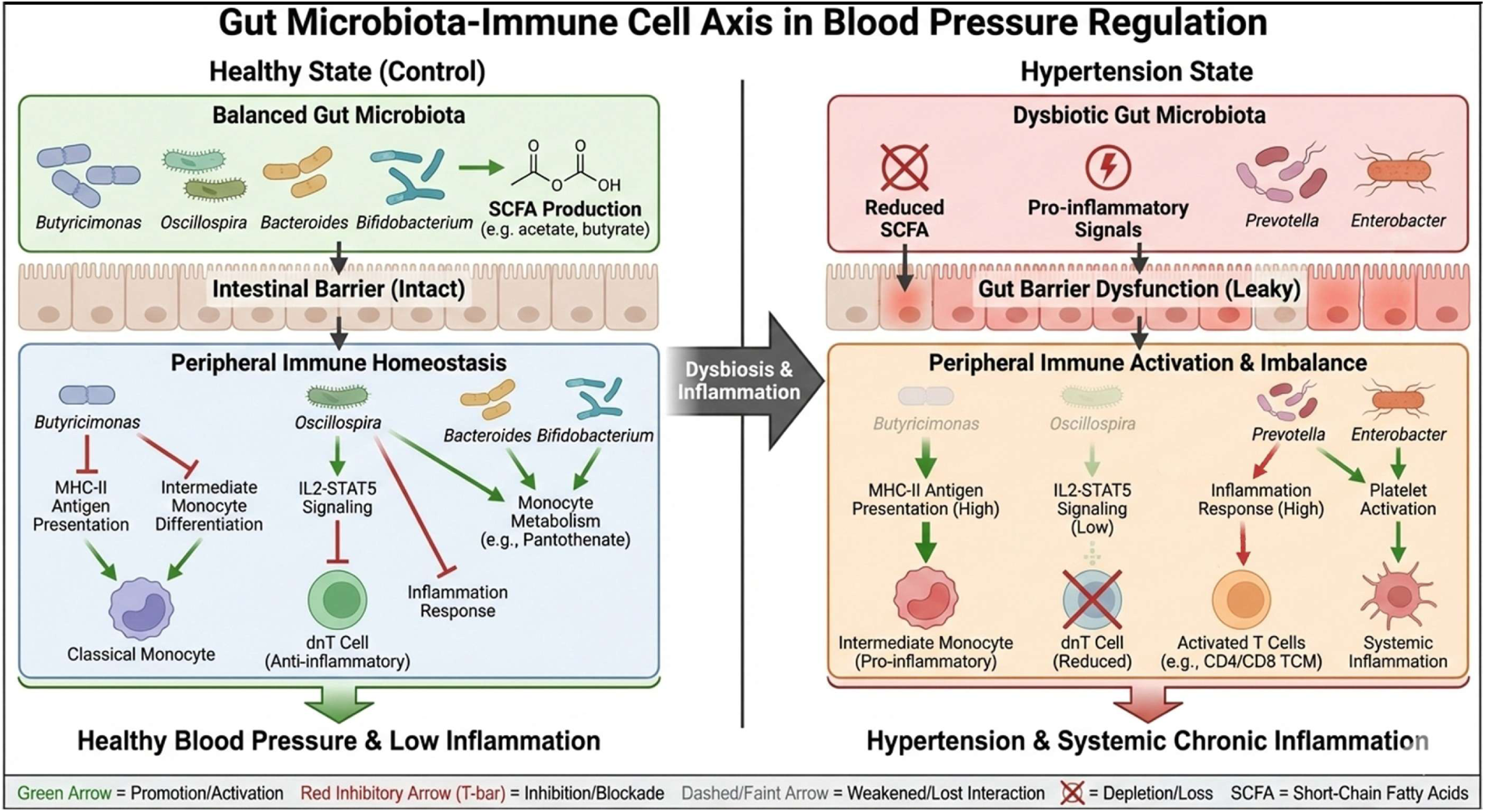
Schematic summary of the main findings and the proposed mechanism

## Acknowledgements

We thank National Natural Science Foundation of China [62102065, 62271353], Joint Funds for the innovation of science and Technology, Fujian province (Grant number: 2022J05055) and Fujian Medical University Research Foundation of Talented Scholars [XRCZX2022003] provided financial support.

## Data availability

Data from the CPH16S cohort are available at https://github.com/SMUJYYXB/GGMP-Regional-variations. Raw sequencing data for the CH16S cohort can be accessed from the NCBI BioProject database under accession number PRJNA417579. Data for the CHMG cohort are available from the curatedMetagenomicData database. Raw single-cell RNA sequencing data are available in the NCBI BioProject database under accession number PRJNA1273480. The data and code used in this study can be accessed in the GitHub repository at https://github.com/as147596/ScMicroBP

## Author Contributions

WL and SH worked on the original draft, handled data, performed analysis, and created visualizations. YZ, SL helped with writing, reviewing, and editing, and also contributed to the methods and supervision. HY, FT and SS managed the project and reviewed and edited the manuscript. FY secured funding, led the project, and contributed to the original draft.

## Corresponding authors

Correspondence to Fenglong Yang

## Consent for publication

All authors have consented to the publication of this work.

## ethics approval statement

This study was based exclusively on publicly available datasets. No human participants or animals were directly involved. Therefore, ethical approval was not required.

## patient consent statement

Not applicable. This study analyzed only publicly available datasets and did not involve direct contact with human participants.

## permission to reproduce material from other sources

Not applicable. No previously published figures or tables were reproduced in this study.

## clinical trial registration

Not applicable. This study did not involve a clinical trial.

## List of abbreviations

CHP16s: Chinese Prospective Hypertension 16S Cohort
CH16s: Colombia Hypertension 16S Dataset
CHMG: Chinese Hypertension Metagenomic Dataset
scBPS: Single-cell Bacterial Polygenic Score
LEfSe: Linear Discriminant Analysis Effect Size
SparCC: Sparse Correlations for Compositional data
RMT: Random Matrix Theory
GWAS: Genome-Wide Association Study
SNP: Single Nucleotide Polymorphism
KEGG: Kyoto Encyclopedia of Genes and Genomes
ATE: Average Treatment Effect

